# Mother’s curse is pervasive across a large mito-nuclear *Drosophila* panel

**DOI:** 10.1101/2020.09.23.308791

**Authors:** Lorcan Carnegie, Max Reuter, Kevin Fowler, Nick Lane, M. Florencia Camus

## Abstract

The maternal inheritance of mitochondrial genomes entails a sex-specific selective sieve, whereby mutations in mitochondrial DNA can only respond to selection acting directly on females. In theory, this enables male-harming mutations to accumulate in mitochondrial genomes if they are neutral, beneficial, or only slightly deleterious to females. Ultimately, this bias could drive the evolution of male-specific mitochondrial mutation loads, an idea known as mother’s curse. Earlier work on this hypothesis has mainly used small *Drosophila* panels, in which naturally-sourced mitochondrial genomes were coupled to an isogenic nuclear background. However, the lack of nuclear genetic variation has precluded robust generalization. Here we test the predictions of mother’s curse using a large *Drosophila* mito-nuclear genetic panel, comprising 9 isogenic nuclear genomes coupled to 9 mitochondrial haplotypes, giving a total of 81 different mito-nuclear genotypes. This enables systematic testing of both mito-nuclear interactions and mitochondrial genetic variance. Following a predictive framework, we performed a screen for wing centroid size, as this trait is highly sexually dimorphic and depends on metabolic function. We confirmed that the trait is sexually dimorphic, and show high levels of mito-nuclear epistasis. Importantly, we report that mitochondrial genetic variance has a greater impact on male versus female *Drosophila*, in 8 out of the 9 nuclear genetic backgrounds. These results demonstrate that the maternal inheritance of mitochondrial DNA does indeed modulate male life-history traits in a more generalisable way than previously envisaged.

## Introduction

Mitochondria are essential organelles for life in eukaryotes, taking centre-stage in the process of cell respiration. Respiration is unusual in that the respiratory complexes within mitochondria are composed of proteins encoded by two different genomes, the nuclear and the mitochondrial. These two genomes must work harmoniously not only to provide cellular energy but also the precursors for most macromolecule synthesis. Consequently, their interaction is vital for the maintenance of mitochondrial integrity and the viability of eukaryote life. Almost without exception, eukaryotes from protists to animals have retained these two genomes (Rand et al. 2004; Lane 2005; Wolff et al. 2014). It was long assumed that purifying selection would remove non-neutral genetic variation within the mtDNA, given the ‘haploid’ nature of this genome and the crucial role they play in energy production (Ballard and Whitlock 2004; Dowling et al. 2008; Cooper et al. 2015). As such, the mitochondrial genome was harnessed as the exemplary molecular marker on which to base evolutionary and population genetic inferences, facilitated by its high mutation rate, maternal inheritance and general lack of recombination (Lynch 1997; Pesole et al. 2000; Saccone et al. 2000; Ballard and Whitlock 2004; Galtier et al. 2009). Over the past two decades, however, a rising number of studies has challenged this assumption of neutrality (Ballard and Kreitman 1994; Rand 2001; Dowling et al. 2008). In particular, numerous studies have revealed that mitochondrial genetic variance sourced from separate populations contributes to the expression of a wide-range of life-history traits (James and Ballard 2003; Maklakov et al. 2006; Melvin and Ballard 2006; Dowling et al. 2007; Wolff et al. 2014; Zhu et al. 2014; Camus et al. 2015; Jelic et al. 2015; Immonen et al. 2016; Salminen et al. 2017).

Mitochondria are maternally inherited in most species, and so natural selection acting on the mitochondrial genome is effective only in females (Rand 2001). Mutations in mtDNA that are beneficial, neutral or even slightly deleterious to males can be selected for in the population, whereas mutations that are detrimental to females should be removed from the population via strong purifying selection (Frank and Hurst 1996; Gemmell et al. 2004). Uniparental inheritance means that males inherit mutations that are selected through the female lineage, even if these mutations are detrimental to them (Innocenti et al. 2011). Through evolutionary time we expect males to accumulate mitochondrial mutation loads consisting of male-biased deleterious mutations. The process leading to sex-biased mutation accumulation has been termed ‘sex-specific selective sieve;’ (Innocenti et al. 2011), or the ‘mother’s curse hypothesis’ (Gemmell et al. 2004).

The idea of the mother’s curse was first described in the 1990s (Frank and Hurst 1996) with further theoretical support proposed the following decade (Gemmell and Allendorf 2001; Gemmell et al. 2004). Efforts have been made to form testable predictions for this hypothesis, with a recent review highlighting a predictive framework to test for the curse (Dowling and Adrian 2019). The first prediction is that not all traits will be equally susceptible to the accumulation of male-biased mitochondrial mutation loads (Friberg and Dowling 2008; Innocenti et al. 2011). In particular, metabolically demanding traits that exhibit sexual dimorphism in expression are most likely to be targets of mother’s curse. This is because the mitochondrial genome underpins most metabolic traits, given the crucial role that mtDNA plays in energy production. When it comes to optimising mitochondrial function for the male homologues of sexually dimorphic traits, males will presumably no longer be able to rely on the female-mediated adaptation of the mtDNA sequence. This is unless nuclear alleles arise that compensate for male-harming mtDNA mutations, which given the difference in evolution rates between mitochondrial and nuclear genes necessarily lags behind (Connallon et al. 2018). In other words, mitochondrial mutations that are metabolically selected for in females may be detrimental to male metabolic demands, given that metabolism is itself a highly dimorphic trait.

Second, if populations harbour mitochondrial genomes comprising male-biased mitochondrial mutation loads, then we should observe greater levels of mitochondrial genetic variance underpinning male relative to female phenotypes. When it comes to the inter-population prediction, mitochondrial haplotypes will evolve along their own population-specific trajectories and accumulate their own distinct pools of male-biased mtDNA mutations with deleterious effects. Purifying selection, however, should remove any such mutations from mtDNA haplotypes that exert deleterious effects on females. These mutations can be unmasked my placing mitochondrial genomes alongside a foreign nuclear genome, where males cannot rely on male-specific coadapted alleles. Thus, when sampling mtDNA haplotypes from distinct populations, we expect greater levels of mitochondrial haplotypic variance underlying the expression of male compared with female phenotypes.

The first conclusive empirical validation of mother’s curse was obtained in a study that examined the effects of mitochondrial variation on genome-wide patterns of nuclear gene expression. Specifically, by placing five different mitochondrial haplotypes alongside an isogenic nuclear background, approximately 10% of the nuclear transcripts were found to be differentially expressed in males relative to females (Innocenti et al. 2011). Interestingly, these differentially expressed transcripts were mostly localised to the male reproductive system (testes, accessory glands, ejaculatory duct), while having no major effect on male non-reproductive or female tissues. More recently, the scope of mother’s curse has broadened beyond reproductive traits (Smith et al. 2010) to other life-history traits including ageing (Camus et al. 2012) and metabolic rate (Nagarajan-Radha et al. 2020).

While support for the mother’s curse hypothesis is growing, one of the limitations of current *Drosophila* work has been unbalanced experimental designs (Rand and Mossman 2020). Studies have either used a single nuclear background coupled to many mitochondrial genomes (Innocenti et al. 2011; Camus et al. 2012; Nagarajan-Radha et al. 2020) or several nuclear genomes coupled to only a few mitochondrial haplotypes (Mossman et al. 2016a; Mossman et al. 2016b). This creates a situation whereby it is difficult to assess how generally the hypothesis might hold across species with high levels of genetic variation. Here we use a new *Drosophila* panel, which comprises a full factorial matrix of 9 worldwide sourced nuclear genomes coupled to 9 mtDNA haplotypes (81 mito-nuclear genotypes in total). Using this panel, we can study both mito-nuclear interactions and mitochondrial genetic variance across several nuclear backgrounds. For each genotype and sex combination we obtain measurements of wing centroid size, which is a highly reliable proxy for *Drosophila* body size (Carreira et al. 2009). Body size is a sexually analogous trait, with clear sexual dimorphism (Partridge et al. 1994; De Jong and Bochdanovits 2003) and high metabolic underpinnings (Guertin and Sabatini 2007; Bryk et al. 2010), making it an ideal candidate trait to test for mother’s curse. Our results show complex interactions between both mitochondrial and nuclear genomes, which modulate wing centroid size. These interactions consistently generate greater variance in males, against all but one nuclear background, confirming that mother’s curse is indeed pervasive across *Drosophila* populations.

## Materials and Methods

### Drosophila stock and maintenance and mito-nuclear panel

The mito-nuclear *Drosophila* panel was produced through the *de novo* full factorial crossing of nine isogenic lines (9 nuDNA x 9 mtDNA). The nine isogenic lines were obtained from the “Global Diversity Panel”, which originate from five different continents and vary in phylogeographic relatedness (Early and Clark 2013; Grenier et al. 2015). Genetic diversity in this panel is representative of that between fruit fly populations world-wide. More specifically they are sourced from: **A**/ZIM184 (Zimbabwe), **B**/B04 (Beijing), **C**/I16 (Ithaca), **D**/I23 (Ithaca), **E**/N14 (Netherlands), **F**/N15 (Netherlands), **G**/T01 (Tasmania), **H**/T23 (Tasmania), **i**/N01 (Netherlands). Using a balancer chromosome crossing scheme (Zhu et al. 2014), we replaced mitochondrial and nuclear genomes. Consequently, the panel contains 81 mito-nuclear genotypes, also known as lines, nine of which are coadapted and 72 are disrupted. Following the balancer chromosome crossing scheme, the complete panel was backcrossed to their respective nuclear genome for 3 generations. We note that one of the mito-nuclear genotypes (F_nuc_×A_mito_) was not viable due to a strong incompatibility, and so for all experiments we proceeded with 80 genotypes.

Lines were propagated by 4-day old parental flies, with approximate densities of 80-100 eggs per vial. Flies were kept at 25 °C and 50% humidity, on a 12:12 hour light:dark cycle, and reared on 8 mL of cornmeal-molasses-agar medium per vial (see **Table S1** for recipe), with *ad libitum* live yeast added to each vial to promote female fecundity. All lines had been cleared of potential bacterial endosymbionts, such as *Wolbachia*, through a tetracycline treatment at the time the lines were created. Clearance was verified using *Wolbachia*-specific PCR primers (ONeill et al. 1992).

### Wing Centroid Size Measure

Focal flies from all genotypes were propagated via two sets of two sequential 4-hour lays. The each lay contained 10-50 flies form each genotype. After second lay oviposition, all vials were cleared. For each lay, vials were density controlled to contain 80-100 eggs. Focal flies were left to develop in their vials for 14 days. At day 14, all flies had emerged and were sexually mature. On this day, flies of each were split by sex and flash-frozen in Eppendorf tubes for subsequent wing analysis.

Fifteen right wings from each sex, and genotype, were pulled and placed on a glass slide using double-sided tape. Along with a 10 mm scale bar, each wing was photographed at 2.5x magnification using an INFINITY Stereo Microscope attached to a Macintosh computer. The programs ‘tpsUtil’ ‘tpsDIG2’ and ‘COORD GEN 8’ were consecutively used to determine the wing centroid size of each photographed wing from eight standard landmarks.

### Statistical Analyses

Our data followed a normal distribution, we used linear models to analyse the data. We first created a model to examine all factors. Centroid size was a response variable with sex, mitochondrial genome and nuclear genome (plus all their interactions) modelled as fixed factors. To further probe this three-way interaction, we divided the dataset by sex and analysed the male and female data separately. For this we used the same model framework as mentioned before; wing centroid size as a response variable with mitochondrial genome, and nuclear genome (plus their interaction) as fixed factors. All models were performed using the *lm/lmer* function in R version 3.3.2 (Team 2016).

To visualise signatures of mother’s curse we standardised all datapoints by their respective sex- and nuclear-specific mean. This way we are able to better compare mitochondrial genetic variance across all nuclear genomes and both sexes (**Figure S1**). To statistically test for mother’s curse predictions, we calculated the mitochondrial coefficient of variation for each nuclear and sex combination using untransformed data. This was done by computing bootstrapped mitochondrial coefficients of variation (CV_m_), in which trait means were resampled with replacement (1000 replicates). This procedure was performed using the *boot* package implemented in R. We used a sign test to test whether male mitochondrial coefficients of variation were overall larger than females and was implemented using the *BDSA* package in R version 3.3.2 (Team 2016).

Given that sequences are available for the protein coding genes of all mitochondrial haplotypes used in our study (Early and Clark 2013), we also tested if there was a correlation between the genetic distance between strains and their phenotypic divergence. To this end, we created matrices of genetic and phenotypic distances between strains (**Table S2**). Genetic distance was quantified as the total number of SNPs difference between lines – excluding the hypervariable region (D-loop). This is because the hypervariable region *Drosophila melanogaster* is a 4.7 kb region with >85% AT-richness; making it very difficult to accurately map reads. Phenotypic matrices were specific to each experimental treatment (performed per sex and nuclear genotype) and phenotypic trait (centroid size). We used a Mantel test for matrix correlation between two dissimilarity matrices, with 10,000 permutations. Mantel test were implemented with the “*mantel*.*rtest*” function from the R package *ade4* (Dray and Dufour 2007)

## Results

### Mito-nuclear epistasis

Our models show sexual dimorphism for wing centroid size, with females having larger centroid size than males (F = 13328.2098, p < 0.001, **Figure 1A, Table S3**). Our full model also revealed effects of both mtDNA and nuDNA, with a significant 3-way interaction between all main factors (mtDNA × nuDNA × sex: F = 1.5639, p = 0.003481). When splitting the models by sex, we find significant mtDNA by nuDNA interactions for both males and females (male: F = 5.1517, p < 0.001; female: F = 4.4962, p < 0.001, **Figure 2, Table S3**). We were interested in the effect size of each genetic component to wing size. For this we calculated the proportion of variance explained by each effect using sums squared derived from the models. Our results show that for both sexes, a large proportion of the variance in wing size is explained by the nuclear genome (female: 51%, male: 49%, **Table S4**). While we found mitochondria to contribute less to the overall variance in wing size observed in our data, we found it to be higher in males (2.3%) than females (0.9%). Similarly, we found the males to have a higher proportion of variance explained by the interaction (15.6%) term than females (10.8%).

**Figure 1:**
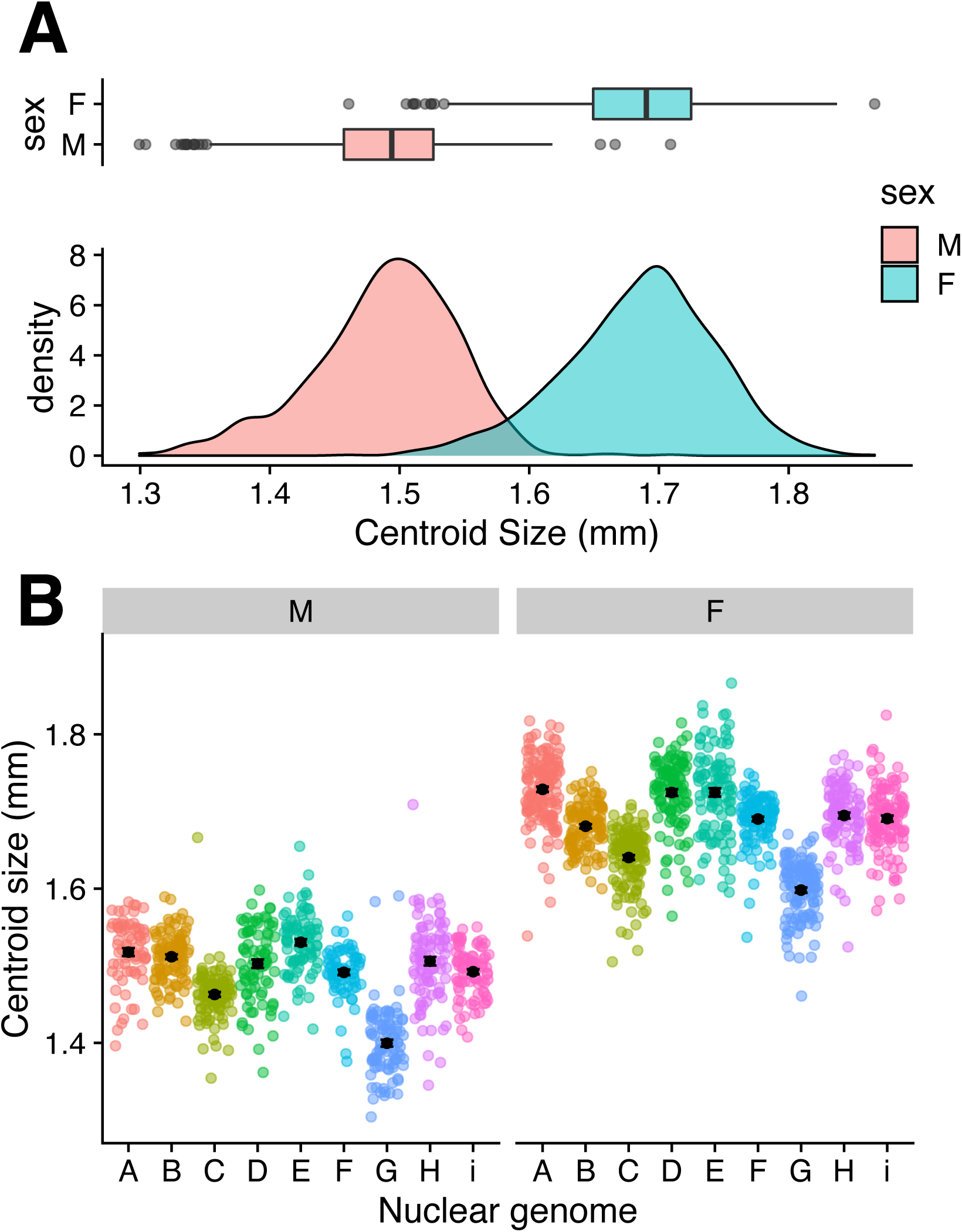
Wing centroid size measurements. (**A**) Broad patterns of sexual dimorphism for this trait with females being larger on average than males. (**B**) patterns of nuclear genetic variation for each sex, showing significant nuclear genetic effects driving centroid size. Individual datapoints are coloured by nuclear genome.

**Figure 2:**
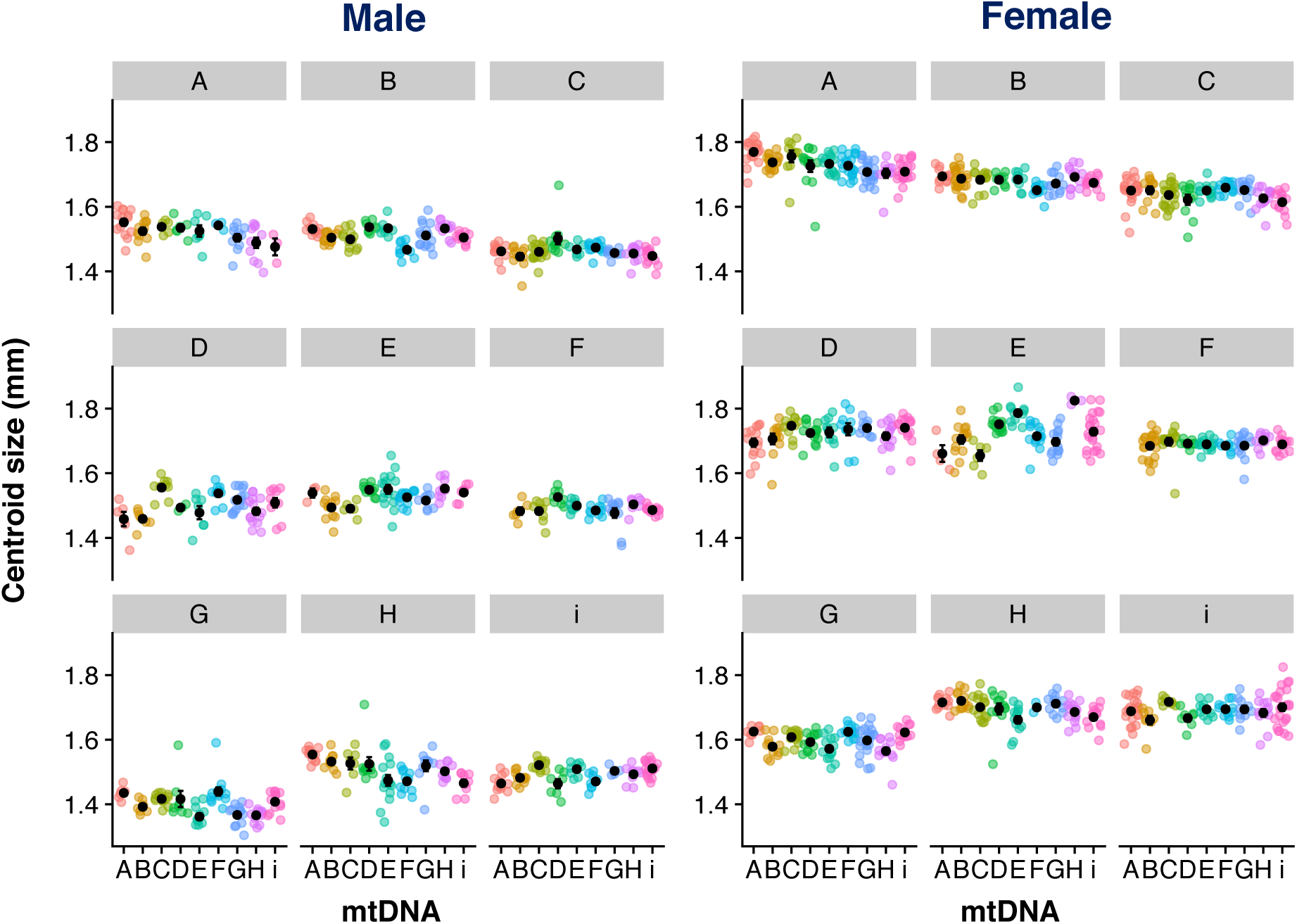
Wing centroid size across all treatments (mean ± SE). Centroid size for males (left) and females (right). Each box corresponds to a specific nuclear genome, with mitochondrial genetic variation shown within each nuclear genome on the X-axis. Individual datapoints are coloured according to the mtDNA genome. Note that the genotype F_nuc_×A_mito_ was not sampled in this experiment as this mito-nuclear combination is completely incompatible.

Overall, there were high levels of genetic correlations between the sexes (r_mf_ = 0.828, p < 0.001, **Figure S2**), indicating that the directionality for each mito-nuclear genotype was concordant between the sexes. Within each nuclear genome we found significant genetic correlations for most nuclear genomes (**Figure S2, Table S5**). We did not find significant genetic correlations for nuclear genomes C, F and I although the directionality of the relationship was positive in all three cases (r_C_ = 0.1652, r_F_ = 0.4303, r_i_ = 0.6449, **Table S5**).

### Testing mother’s curse predictions

An important prediction of mother’s curse is that it should result in mitochondrial genomes that harbour mutation loads that are more pronounced in males and that these loads can be uncovered by demonstrating greater levels of functional mitochondrial genetic variance in males than in females. To this end, we calculated coefficients of variation for each nuclear genome and sex combination, and our results show that overall males have significantly higher levels of mitochondrial genetic variance than females (p = 0.03906 [0.0015, 0.0097], **Figure 3, Table S6**). Upon closer inspection, this rule is true for all but one nuclear genome (“E” nuclear genome). Moreover, we find that for 6 of the 8 genotypes that have greater mitochondrial coefficients of variation in males even raw variance is higher—despite males being smaller (**Table S6**).

**Figure 3:**
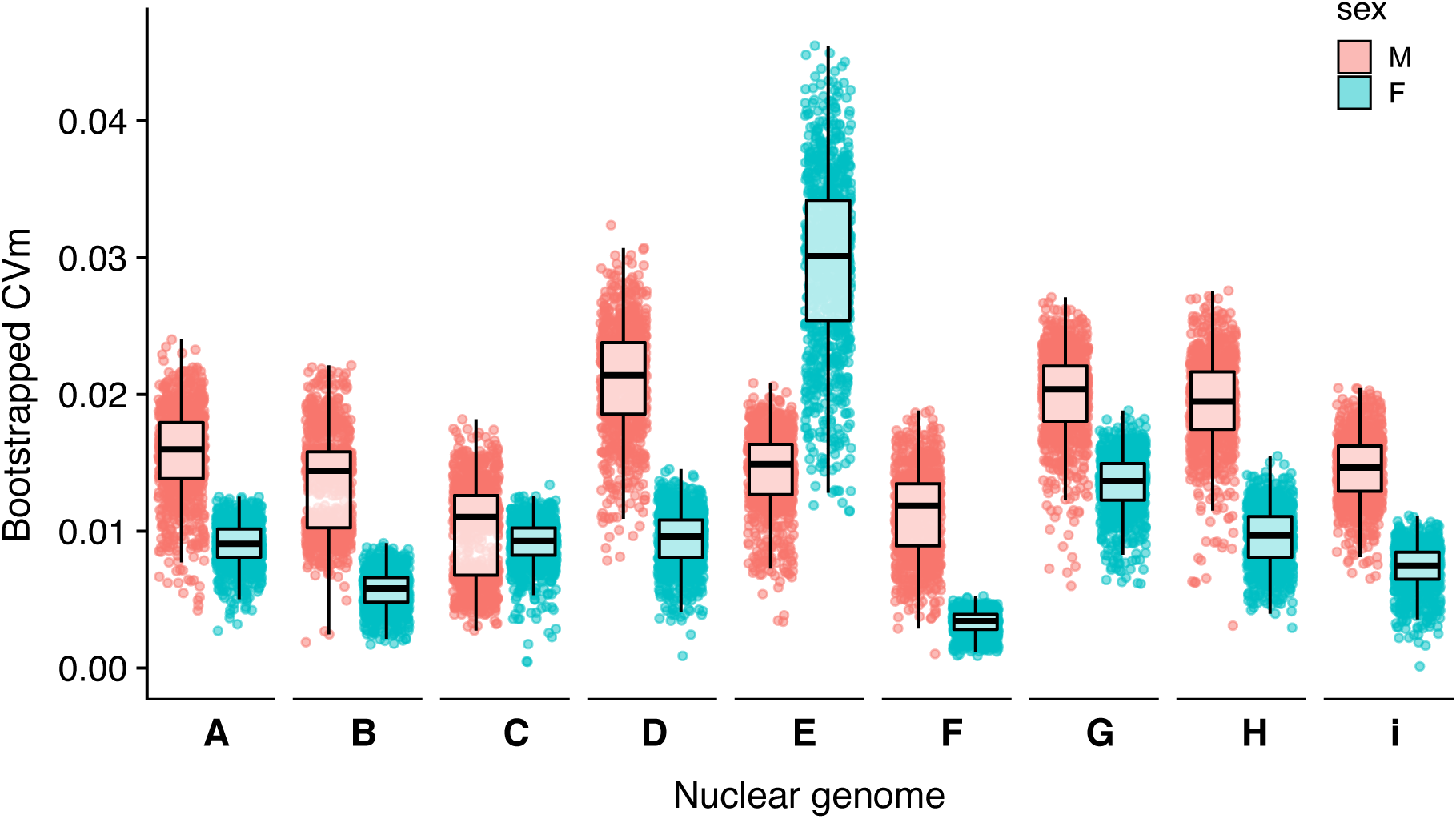
Mitochondrial genetic variance. Bootstrapped mitochondrial coefficients of variation within each nuclear genome for both sexes.

### Relationship between mitochondrial genetic distance and phenotypic differentiation

We asked the question of whether the degree of coadaptation influenced phenotypic differentiation. For this we used Mantel tests to correlate two matrices: one comprising the mitochondrial genetic differences between the haplotypes (total SNPs – **Table S3**), with the other consisting of the mean phenotypic divergence from the coevolved strain. This analysis was performed for each nuclear genome separately. Given that we were unable to collect phenotypic data for one mito-nuclear genotype (due to a severe incompatibility), the analysis for the F nuclear background was run using 8 eight genotypes rather than nine. This analysis shows that there is no relationship between genetic and phenotypic divergence for females (**Figure 4**). Males showed similar results to females, however, for 2 out of the 9 nuclear genomes there was a significant positive association between genotype and phenotype divergence. More specifically, these were nuclear genome A (r = 0.543 p = 0.00391) and H (r = 0.228, p = 0.0158). Such weak or absent correlation between mito-nuclear genotype and phenotype has been reported before (Camus et al, 2020) and reflects the stochastic nature of small numbers of interactions between closely related populations, combined with the requirement for metabolic plasticity between tissues and sexes, which can buffer the effects of mutations.

**Figure 4:**
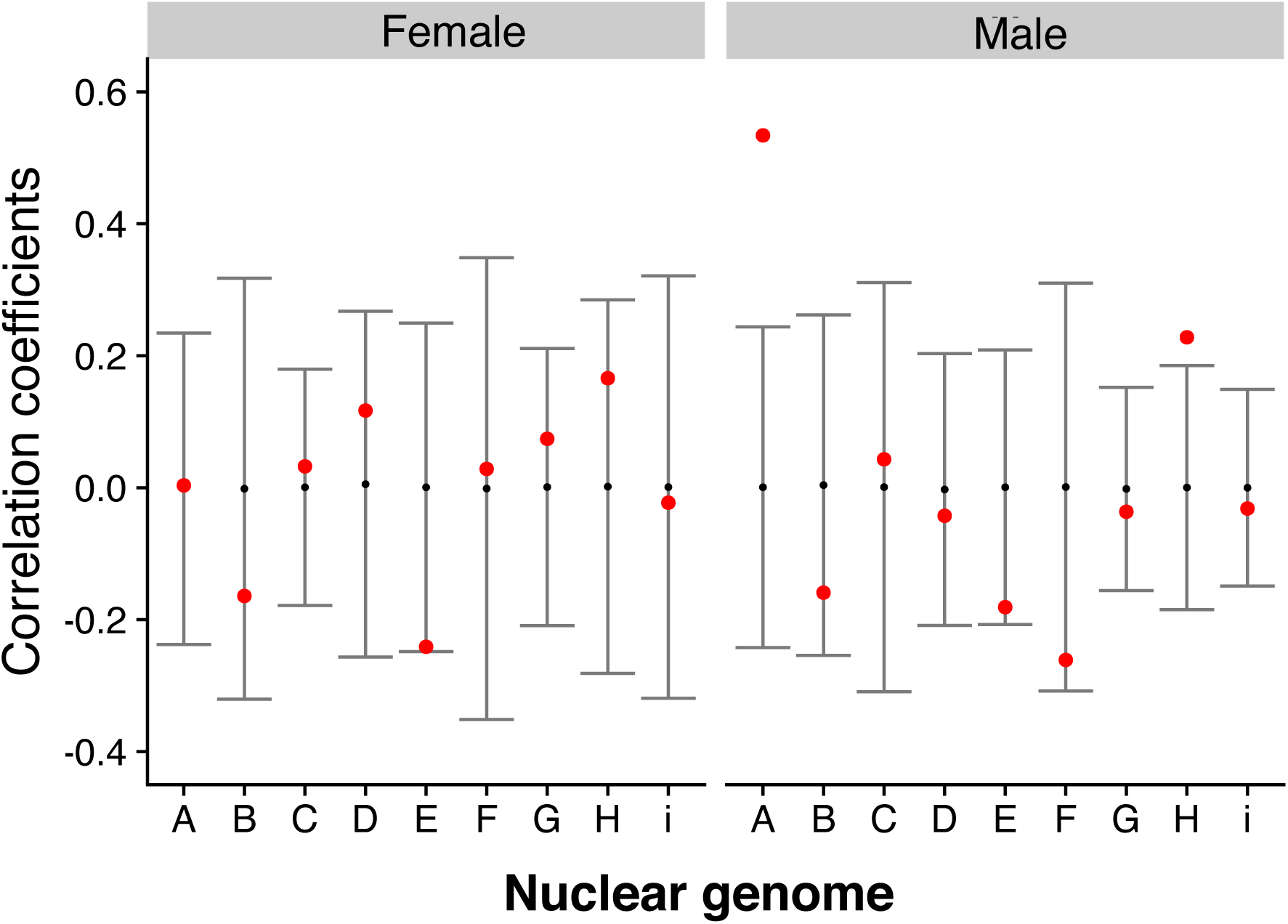
Association between genotypic and phenotypic divergence. Mantel test correlation of two matrices showing the null distribution of the permutations in grey with the mean correlation value for each test in red. Significant associations are when the correlation statistic falls outside of the null distribution. Mitochondrial genetic differences were calculated as total number of SNPs difference (not including the hypervariable region) between all the strains. For the phenotypic matrix, each nuclear genome was considered separately, creating a matrix of the average size difference between the coevolved strain and all other eight strains. Note that for the “F” nuclear genome this analysis was performed with 8 genomes as one of our genotypes was completely incompatible.

## Discussion

The maternal inheritance of the mtDNA causes selection to fix mutations that are neutral or beneficial in females, even if these mutations are harmful in males. Ultimately, this evolutionary process has the potential to drive the accumulation of male-specific mitochondrial mutation loads, a hypothesis known as mother’s curse. While there is some experimental evidence for this hypothesis, one of the most pertinent criticisms is that the limited use of nuclear genetic backgrounds raises the possibility that mothers curse is not generalisable (Rand and Mossman 2020). Here we aim to address this criticism by using a new panel of *Drosophila melanogaster* comprising nine nuclear backgrounds, each coupled to nine mtDNA haplotypes, resulting in 81 new mito-nuclear genotypes. In line with the predictions of mother’s curse (Dowling and Adrian 2019), we measured wing centroid size because the trait is sexually dimorphic and its development exerts high metabolic demands. We find complex interactions between all three factors (sex × nuDNA × mtDNA), indicating important epistatic effects in our dataset. Moreover, we find that mtDNA produces greater variance in males than in females across 8 out of the 9 nuclear backgrounds used in this study. Our findings thus demonstrate that the mother’s curse is more generalisable than previously appreciated.

We first confirmed the sexual dimorphic patterns for wing centroid size, a pattern that has been observed in many earlier studies (Carreira et al. 2009; Gidaszewski et al. 2009). We also observed high levels of nuclear genetic variance within each sex for this trait, with this variation being mostly sexually concordant. Previous work that considered the contribution of nuclear genetic variance for wing size/shape using the DGRP strains again found very similar results (Pitchers et al. 2019). Our analyses demonstrate that a large proportion of this variation can be explained by the nuclear genome. Given that the nuclear backgrounds used in this study are representative of five different worldwide populations, we could have investigated whether this variation is due to the different geographical regions; however our sample sizes per location were not adequate. During the creation of the panel, we selected fly strains that maximised both nuclear and mitochondrial genetic diversity, so it is possible that we captured many rare genomes from each population. The fact that each nuclear genome is inbred also complicates population-specific comparisons, so we deemed it best to treat them as independent genetic units.

Not all studies that looked for signatures of Mothers Curse have found male-biased effects (Mossman et al. 2016a; Mossman et al. 2016b; Dordevic et al. 2017; Mossman et al. 2017). For example, mixed results have been obtained when examining mitochondrial genetic effects on nuclear gene expression. More specifically, while Innocenti et al. (Innocenti et al. 2011) found male-biased genes being affected by replacing the mtDNA genome, work by Mossman et al. (Mossman et al. 2016b) did not find these patterns. There are many differences between the two studies, with one of the main differences being that the latter study used mito-nuclear flies which combined genomes from different species (*D. melanogaster* and *D. simulans*) and limited the number of mtDNA genomes sampled. Recent efforts have been made to create robust experimental guidelines to systematically test the predictions of mother’s curse (Dowling and Adrian 2019). One of the main guidelines specifies that adequate levels of mitochondrial genetic variation be tested, as small numbers of mitochondrial genomes can easily under or over-represent variance. With this in mind, when creating our mito-nuclear panel for our study, we aimed to maximise the number of mtDNA genomes. While we do not have as many mtDNA genomes as some previous studies (Camus et al. 2012; Camus et al. 2015), we have captured far more nuclear genetic variation, enabling us to test for mother’s curse across multiple nuclear environments.

Our results support the mother’s curse hypothesis in two ways. First, the variance accounted for by mtDNA in our linear models is greater in males than females. While the contribution of the mitochondrial genome to wing size variation is clearly much less than the nuclear genome, we also see an equivalent pattern when looking at the mito × nuclear interaction term. Previous work has found similar results when considering the contribution of the two genomes on the transcriptome, with the nuclear genome having a much greater effect than mtDNA (Mossman et al. 2016b; Mossman et al. 2017). We then measured the mitochondrial coefficients of variation and found similar patterns, whereby for most nuclear genomes, males showed higher levels of variation than females (**Figure 3**). The only nuclear background where this was not the case was “E” (strain N14), where changing the mtDNA genome had a large effect on females. While we do not know the exact mechanisms driving this response, we note that this nuclear background was particularly sick during rearing. We also noticed that mtDNA replacement for this nuclear background tends to decrease wing size, suggesting that the “E” nuclear background could particularly susceptible to changes in physiology (**Table S7**). In sum, our experimental design was formulated taking into account the framework proposed to test the predictions of mother’s curse: we believe we have generated a robust dataset in favour of the hypothesis.

Given our experimental design, we were also able to investigate predictions for mito-nucelar coadaptation, whereby coadapted genomes outperform disrupted combinations. We therefore examined whether phenotypic divergence from the coadapted mito-nuclear combination correlated with genetic divergence. In other words, does greater genetic disruption result in larger phenotypic differences? While we could see no association in females, we did find a significant positive association in two out of the nine nuclear genomes in males. These results suggest that mito-nuclear interactions can exert significant, albeit inconsistent, effects on life-history traits in different contexts. Previous work has likewise reported inconsistent results when looking at mitochondrial genetic relationships with fecundity and longevity (Camus et al. 2012; Camus et al. 2020). It could be that the effect and magnitude of different mtDNA SNPs depends on the environment that the nuclear background produces, so a SNP with a small effect in one nuclear background could have a large effect in another. Relatedly, the functional nuclear background varies from tissue to tissue and between sexes, reflecting differences in gene expression and metabolic demands. This means that the mitochondrial genome must work effectively across multiple backgrounds, which likely limits the cumulative impact of SNPs, undermining correlations with genetic distance (Camus et al. 2020). Another caveat when considering coadaptation using this trait is that we are assuming that a larger wing reflects higher fitness. While this tends to be true in females, it is less clear with males (Lefranc and Bundgaard 2000). That is because optimal male body size appears to be under stabilising selection (Lefranc and Bundgaard 2000), with larger males more likely to harm to females (Pitnick and Garcia-Gonzalez 2002). So while there is a lot of evidence that metabolic deficits result in smaller flies (Bryk et al. 2010), there might also be an upper fitness optimum to how large *Drosophila* can get.

In conclusion, we provide robust evidence for the mother’s curse hypothesis. Our experimental design aimed to address some of the weaknesses of earlier work, notably the use of single nuclear backgrounds. We report greater levels of mitochondrial genetic variance affecting male traits in all but one of the nuclear backgrounds. Future work using this *Drosophila* panel will investigate other phenotypic traits to examine mother’s curse under a multivariate framework.

## Acknowledgments

This work was funded by the Leverhulme Trust Grant [RPG-2019-109] to MFC/KF/MR/NL. We thank Rebecca Finlay for help with running experiments

## Author Contributions

MFC, KF, and MR created the mito-nuclear *Drosophila* panel. LC ran the experiments, collected all data, and performed preliminary analyses. MFC designed the experiment, assisted with data collection and performed analysis. MR helped with statistical analyses. MFC lead in manuscript writing with helpful contributions from NL.

## Data Accessibility

All data will be made available on DRYAD repository upon acceptance.

## Notes

### Competing Interest Statement

The authors have declared no competing interest.

