## Supplementary for "Mother’s curse is pervasive across a large mito-nuclear *Drosophila* panel"

**Table S1:** Drosophila rearing medium recipe. This recipe makes for 2L worth of food.

| Ingredient | Quantity |
|---|---|
| Yeast | 200g |
| Sugar | 200g |
| Agar | 30g |
| Water | 2L |
| Propionic acid (preservative) | 6mL |
| Nipagin (preservative) | 60mL |

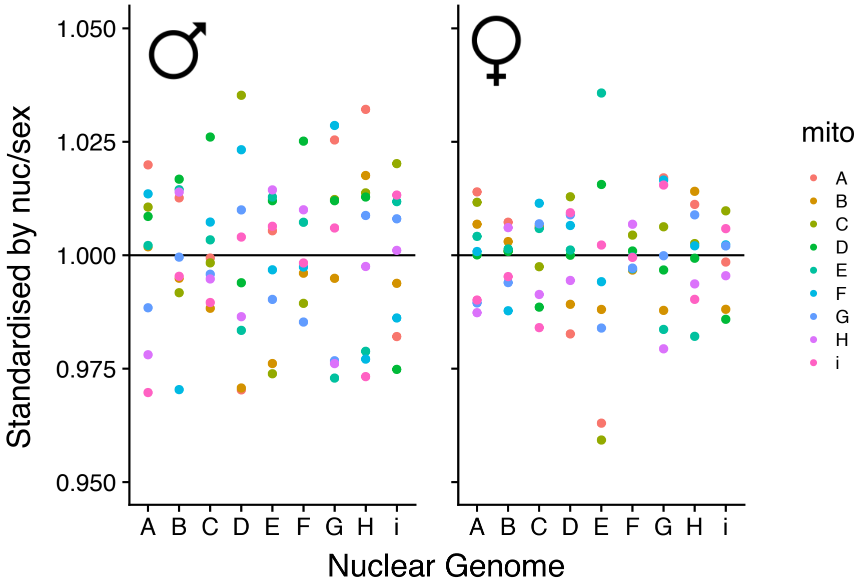


**Figure S1:** Mitochondrial genetic variation for each nuclear genome and sex combination. Values have been standardised by the mean nuclear genome and sex-specific mean and allows for better visualisation of the patterns observed.

**Table S2:** Mitochondrial genetic differentiation matrix. Below the diagonal we show the total number of SNPs difference between the mtDNA genomes, with the top of the diagonal showing the non-synonymous SNPs.

| | **A** | **B** | **C** | **D** | **E** | **F** | **G** | **H** | **i** |
|---|---|---|---|---|---|---|---|---|---|
| **A** | - | 1 | 0 | 1 | 1 | 0 | 0 | 0 | 4 |
| **B** | 32 | - | 1 | 2 | 2 | 1 | 1 | 1 | 5 |
| **C** | 27 | 35 | - | 1 | 1 | 0 | 0 | 0 | 4 |
| **D** | 27 | 35 | 8 | - | 2 | 1 | 1 | 1 | 5 |
| **E** | 25 | 31 | 22 | 22 | - | 1 | 1 | 1 | 5 |
| **F** | 27 | 37 | 8 | 12 | 24 | - | 0 | 0 | 4 |
| **G** | 29 | 31 | 20 | 24 | 14 | 24 | - | 0 | 4 |
| **H** | 24 | 26 | 25 | 25 | 13 | 27 | 9 | - | 4 |
| **i** | 62 | 65 | 56 | 62 | 58 | 58 | 56 | 61 | - |

**Table S3:** Outputs from linear models where centroid size was modelled with mtDNA, nuDNA and sex as fixed factors.

**Full Model**

| | Df | Sum Sq | Mean Sq | F value | Pr(>F) |
|---|---|---|---|---|---|
| mito | 8 | 0.17 | 0.0213 | 16.7597 | < 0.001 |
| nuc | 8 | 3.361 | 0.4201 | 331.2966 | < 0.001 |
| sex | 1 | 16.9016 | 16.9016 | 13328.2098 | < 0.001 |
| mito × nuc | 63 | 0.6089 | 0.0097 | 7.621 | < 0.001 |
| mito × sex | 8 | 0.0235 | 0.0029 | 2.3153 | 0.01806 |
| nuc × sex | 8 | 0.1194 | 0.0149 | 11.7717 | < 0.001 |
| mito × nuc × sex | 63 | 0.1249 | 0.002 | 1.5639 | 0.003481 |
| Residuals | 1688 | 2.1406 | 0.0013 | | |

**Female**

| | Df | Sum Sq | Mean Sq | F value | Pr(>F) |
|---|---|---|---|---|---|
| mito | 8 | 0.03279 | 0.004099 | 3.0345 | 0.00224 |
| nuc | 8 | 1.80344 | 0.22543 | 166.8931 | < 0.001 |
| mito × nuc | 63 | 0.38261 | 0.006073 | 4.4962 | < 0.001 |
| Residuals | 977 | 1.31968 | 0.001351 | | |

**Male**

| | Df | Sum Sq | Mean Sq | F value | Pr(>F) |
|---|---|---|---|---|---|
| mito | 8 | 0.055 | 0.006875 | 5.9547 | < 0.001 |
| nuc | 8 | 1.14963 | 0.143704 | 124.4664 | < 0.001 |
| mito × nuc | 63 | 0.37472 | 0.005948 | 5.1517 | < 0.001 |
| Residuals | 711 | 0.82089 | 0.001155 | | |

**Table S4:** Proportion of variance explained by explanatory variables for both male and female models.

| | male | female |
|---|---|---|
| mito | 0.023 (2.3%) | 0.009 (0.9%) |
| nuc | 0.479 (47.9%) | 0.510 (51%) |
| mito × nuc | 0.156 (15.6%) | 0.108 (10.8%) |

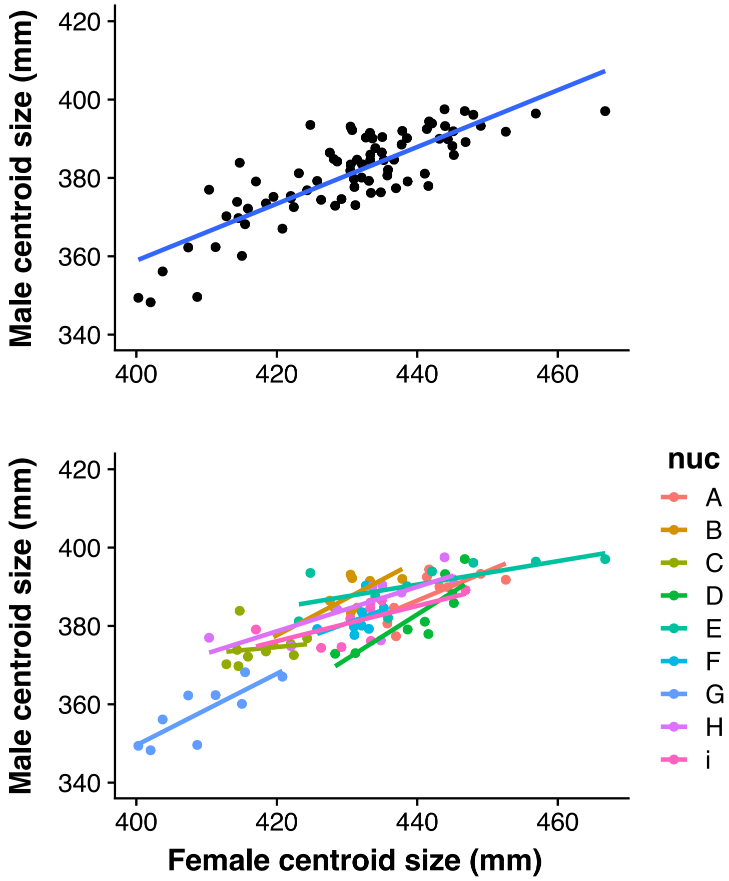


**Figure S2: Centroid size correlations.** We performed correlations for centroid size between males and females (top). The bottom plot shows the same correlation, but distinct nuclear genomes are coloured, for better visualization of mitochondrial genetic correlations.

**Table S5:** Mitochondrial genetic correlations per nuclear genome

| nuc | r | p | CI | |
|---|---|---|---|---|
| A | 0.7131035 | **0.03103** | 0.09304822 | 0.93460699 |
| B | 0.8026384 | **0.009216** | 0.2966411 | 0.9567597 |
| C | 0.1652414 | 0.6709 | -0.5603762 | 0.7473488 |
| D | 0.8558237 | **0.003245** | 0.4441361 | 0.9691236 |
| E | 0.7026195 | **0.03479** | 0.07217775 | 0.93189557 |
| F | 0.4303116 | 0.2872 | -0.393761 | 0.8709019 |
| G | 0.8172228 | **0.007153** | 0.3348195 | 0.9602065 |
| H | 0.7701874 | **0.01517** | 0.2171246 | 0.9489321 |
| i | 0.6448916 | 0.06075 | -0.0336358 | 0.91649212 |

**Table S6:** Coefficient of variation (CV), variance and standard deviation estimates for both sexes and within each nuclear genome. Shown in bold are cases where the estimates are greater in males than females.

| | CV | | Variance | | Standard deviation | |
|---|---|---|---|---|---|---|
| nuc | male | female | male | female | male | female |
| A | **0.01564599** | 0.00905772 | **0.00200078** | 0.00163517 | **0.04473007** | 0.04043718 |
| B | **0.01405321** | 0.00576693 | **0.00096923** | 0.00084238 | **0.03113245** | 0.02902377 |
| C | **0.01000264** | 0.00904443 | 0.00105992 | 0.00134211 | 0.03255647 | 0.03663489 |
| D | **0.02089639** | 0.00933088 | **0.00236049** | 0.00201607 | **0.04858486** | 0.04490061 |
| E | 0.01441906 | 0.02962826 | 0.00148466 | 0.00309969 | 0.03853125 | 0.05567481 |
| F | **0.01169987** | 0.00335646 | **0.00101766** | 0.00096331 | **0.03190078** | 0.03103725 |
| G | **0.01989798** | 0.01363829 | **0.00199818** | 0.00138559 | **0.04470103** | 0.03722357 |
| H | **0.01915078** | 0.00956285 | **0.00285628** | 0.0014458 | **0.05344417** | 0.03802367 |
| i | **0.0146611** | 0.00726518 | 0.00103125 | 0.00171638 | 0.03211301 | 0.04142919 |

Mito-nuclear coadaptation

One of the questions we wanted answered was if coevolved genotypes had consistently larger wings than disrupted. This is under the assumption that disrupting mito-nuclear interactions would have energetic consequences and flies would be smaller. For this we calculated the differences between the average coevolved wing size and the average disrupted wing size. While we compress a lot of datapoints into a single value, it was the best strategy (to our knowledge) to avoid comparing groups with unequal sample sizes (there are 8-times more disrupted lines than coevolved).

**Table S7:** Degree of coadaptation for each nuclear genome

| nuc | male | female |
|---|---|---|
| A | **0.0377** | **0.027** |
| B | -0.0088 | **0.006** |
| C | -0.0029 | -0.005 |
| D | -0.0095 | 0.000 |
| E | **0.0241** | **0.069** |
| F | -0.0045 | -0.005 |
| G | -0.0371 | 0.000 |
| H | -0.0042 | -0.012 |
| i | **0.0247** | **0.013** |
